# Avoiding accuracy-limiting pitfalls in the study of protein-ligand interactions with isothermal titration calorimetry

**DOI:** 10.1101/023796

**Authors:** Sarah E. Boyce, Joel Tellinghuisen, John D. Chodera

**Affiliations:** Schrödinger, 120 West 45th St, New York, NY 10036; Department of Chemistry, Vanderbilt University, Nashville, TN 37235; Computational Biology Program, Sloan Kettering Institute, Memorial Sloan Kettering Cancer Center, New York, NY

**Keywords:** isothermal titration calorimetry (ITC), propagation of error, entropy-enthalpy compensation

## Abstract

Isothermal titration calorimetry (ITC) can yield precise (±3%) estimates of the thermodynamic parameters describing biomolecular association (affinity, enthalpy, and entropy), making it an indispensable tool for biochemistry and drug discovery. Surprisingly, interlaboratory comparisons suggest that errors of ∽ 20% are common and widely underreported. Here, we show how to reduce precision- and accuracy-limiting errors while obtaining good estimates and minimizing material and time consumed by an experiment. We provide a simple spreadsheet that allows practitioners to identify precision-limiting operations during protocol design, track precision during the experiment, and propagate error to yield realistic final uncertainties.

---

Isothermal titration calorimetry (ITC) [1] is a popular technique for probing phenomena of biological interest, including protein-ligand interactions. While the method consumes more reagents than optical or spectroscopic techniques, it does not require specific labeling of the system under study, and a single experiment can yield estimates of all thermodynamic parameters characterizing a reaction—the association constant *K*_*a*_ (and hence also the standard binding free energy Δ*G* ^*°*^ = − *RT* ln[*K*_*a*_*C °*]^1^), the standard enthalpy change Δ*H* ^*°*^, and the reaction standard entropy Δ*S* ^*°*^. With careful work on a well-behaved system, relative standard errors (RSEs) of 1-3% are regularly achievable [2]. However, in a large-scale survey (ABRF-MIRG’02) in which 14 core ITC facilities studied the association of carboxybenzenesulfonamide (CBS) with bovine carbonic anhydrase II (CAII), the variation among reported binding constants and enthalpies was more than an order of magnitude larger than the standard errors reported by the participants [3]. This unexpectedly large variation has been attributed mainly to errors in titrant (syringe reagent) concentration, which is treated as exact in standard analysis procedures [4]. Failure to propagate these errors into reported results can lead practitioners astray in the interpretation of their data,especially if differences in Δ*G*° or Δ*H* ^*°*^ are of interest—for example, within a structure-activity relationship (SAR) series or in interpretation of enthalpic (Δ*H* ^*°*^) and entropic (− *T* Δ*S* ^*°*^) contributions to binding [5, 6].

We therefore strongly advocate that practitioners report the method by which the titrant is prepared, the uncertainty in titrant concentration, and the resulting total error in thermodynamic parameters including titrant uncertainty in all reports of calorimetric measurements. Otherwise, it *must* be assumed that the reported *K*_*a*_, Δ*H* ^*°*^, and ΔS° are contaminated by errors up to 20%, the best estimate of this unreported error available to date [4]. To aid practitioners in reducing and reporting error, we discuss accuracy-limiting steps in solution preparation and provide a spreadsheet^2^ for automatically tracking uncertainties and propagating their contributions to produce realistic error estimates.

## Error propagation

The general rule for random error propagation for a quantity *f* (*x*, *y*, *z*,…) dependent on *independent* measurements *x*, *y*, *z*,… gives squared standard error *s*_*f*_ in *f* as [7],

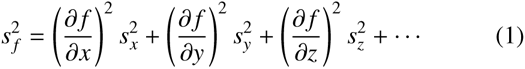

where *s*_*z*_ are the standard errors of the corresponding measurements. This form, based on a Taylor expansion of the function *f*, can be extended to any number of contributing terms (e.g., multiple solution preparation steps) [7]. If *f* can be written *f* (*x*, *y*, *z*,…) = *x*^*i*^*y*^*j*^*z*^*k*^—where *i*, *j*, and *k* are powers to which *x*, *y*, and *z* are raised—then Eq. 1 assumes the simple form for the relative error (*s*_*f*_ |*f*),

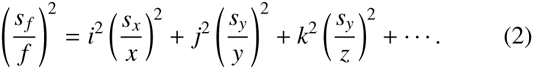

Often, a single term in Eq. 2 will dominate, and the relative error is essentially identical to this contribution. For example, in the calculation of our titrant concentration from mass *m* and volume *v*, *c* = *m*/*v* with (*s*_*m*_|*m*) = 1% and (*s*_*v*_/*v*) = 0.2%, then the s (*s*_*c*_/*c*) ≈ 1%. We will utilize this scheme to propagate error throughout our experiment, as well as to incorporate these errors alongside the least squares fit error in thermodynamic parameters produced by standard calorimetry analysis software. To simplify this process for typical applications, the provided spreadsheet performs much of this error propagation automatically.

## Illustrative application to CAII:CBS

For illustration, we consider the target reaction from the ABRF-MIRG’02 survey [3], the 1:1 association of CBS and bovine CAII, which can be written,

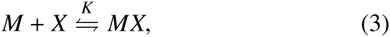

where *M* denotes macromolecule and *X* ligand. This reaction has a *K*_*a*_ ∼ 10^6^M and Δ*H* ∽ -10 kcal/mol [3, 4].

As both protein and ligand may be precious, there is a desire to minimize material use in protein-ligand studies. Using concentrations only as large as necessary also minimizes the need for buffer additives such as DMSO to enhance solubility, reducing agents to prevent crosslinking, and detergents to inhibit aggregation. These additives pose additional experimental challenges, as calorimetrically-measured heats can be sensitive to even small composition mismatches between cell and syringe solutions. Minimizing these effects requires dialysis of the macromolecule by buffer followed by preparation of the ligand in the dialysate. If the ligand is already in solution (e.g. in DMSO stocks), it may not be possible to fully eliminate excipients, leading to potential heat effects due to buffer mismatch even if attempts are made to match compositions.

## Experimental design

In the ABRF-MIRG’02 survey [3], participants employed titrand (cell reagent) concentrations [M]_0_ in the range 7–71 *μ*M. We used an ITC protocol design program [8], which indicated ~3% relative standard error (RSE) in *K*_*a*_ and ∼ 1% for Δ*H* ^*°*^ was possible with our instrument (a GE/MicroCal VP-ITC) using [M]_0_ = 10 *μ*M (consuming ~0.5 mg protein per experiment). While this gives *c* = *K*[*M*]_0_ ≈ 10, a key ITC parameter [1], in the low range of the generally recommended 1 ≤ *c ≤* 1000 range [1], high measurement precision can still be obtained at this *c* value by titrating to an optimal titrant:titrand ratio *R*_*m*_ given by,

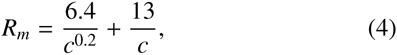

a heuristic expression^3^ obtained from a comprehensive study of precision as a function of *R*_*m*_ [9]. The suggested *R*_*m*_ = 5.3 is significantly greater than the *R*_*m*_ = 2 that is widely used in standard protocols for ITC; with decreasing *c*, use of *R*_*m*_ = 2 progressively limits the fractional conversion of M to MX and thus limits the precision of estimation for both *K*_*a*_ and Δ*H* ^*°*^ [8].

In the present case, use of *R*_*m*_ = 2 would cause significant precision loss, almost doubling the achievable RSEs for *K*_*a*_ and Δ*H*^*°*^.

The same optimization study [9] demonstrated that the experimental precision depended only weakly on the number of injections *m*, recommending *m* = 10 for processes confidently known to involve 1:1 complexation. This is in sharp contrast to ∼ 30 injections often recommended by standard protocols in order to visualize a full sigmoidal (S-shaped) curve in the enthalpogram, which unnecessarily limits precision by reducing the heat per injection (increasing RSEs to 19% and 4%, respectively), as well as increasing the duration of the titration experiment nearly three-fold [9]. Using 10 injections, each of volume *v* = 10 *μ*L, we can compute the approximate syringe concentration [*X*]_*s*_ [10]),

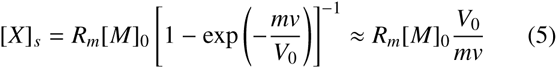

where *V*_0_ is the cell active volume (∼ 1.4 mL for the VP-ITC) and the approximate equality follows if the total titrant injected is small compared to the cell volume (*mv* ≪ *V*_0_). For our experiment, Eq. 5 suggests we should use a purity-corrected titrant concentration [*X*]_*s*_ ∼ 720 *μ*M.

## Syringe backlash and the first-injection anomaly

Our GE/MicroCal VP-ITC instrument has a syringe assembly that utilizes a worm gear which, after the recommended purge-refill process, will cause a titrant shortfall in the first injection unless a “down syringe” command is issued prior to loading the syringe into the sample cell [11]; we therefore executed a 10 *μ*L “down syringe” command immediately after the purge-refill cycle. Because the instrument can take a substantial (but variable) period of time to stabilize at the desired experimental temperature after loading the syringe, significant (> 0.1 *μ*L) diffusive loss can also contribute to a first injection shortfall. We therefore programmed an initial 1 *μ*L “throwaway injection” to avoid the need to correct for diffusive titrant loss during the first 10 *μ*L injection. The contribution from this initial 1 *μ*L “throw-away” injection was excluded from the fitting procedure during analysis. Note that even though we exclude this heat from the analysis, we still need the syringe “down” command to ensure that the correct amount of titrant enters the cell.

## Titrand preparation

The titrand solution, bovine CAII (Sigma-Aldrich, cat no. C2522, ∼30 kDa, Lot No. 071M6261) in PBS buffer, was prepared following the assay conditions out-lined by Myszka et al. [3]. Briefly, the contents of the glass vial containing ~ 5 mg of lyophilized CAII were resuspended in 750 *μ*L filtered buffer and dialyzed overnight in 1 L buffer using a Novagen D-Tube Dialyzer MWCO 3.5 kDa (Cat No. 71506-3, Lot D00131446). The recovered protein was spun for 30 min at 16 300 RPM with no visible precipitate observed. The dialysate was filtered again and used to prepare both titrant and titrand to minimize buffer mismatch heats during the ITC experiment.

The protein concentration was determined spectrophotometrically via absorbance at 280 nm on a NanoDrop ND-1000. The NanoDrop (and similar instruments) utilize small sample volumes (3 *μ*L was used here) and dynamic selection of among path lengths (between 0.2 mm and 1 mm for the ND-1000) to facilitate direct determination of typical protein concentrations without dilution. Here the measured absorbance of 1.18±0.02 at 1 mm path [henceforth written 1.18(2)] length yielded a protein concentration of 235 ± 4 *μ*M using the known molar absorptivity *∊*_280 nm_ = 50070(25) M^−1^ cm^−1^ [3]. The sample was then diluted to [M]_0_ ≈ 10 *μ*M using the purity-corrected post-dialysis concentration. Note that high precision is not generally required for protein concentration determination unless the binding stoichiometry is unknown, as the site parameter *n* absorbs errors in [M]_0_ and *V*_0_ in standard least-squares data analysis [8].

## Titrant preparation

In contrast to titrand preparation, care must be taken to minimize inaccuracies in preparing titrant solutions, because the standard data analysis algorithms treat [X]_*s*_ as exactly known. Thus, a 1% error in [X]_*s*_ produces 1% errors in the estimates of *K*_*a*_ and Δ*H*° [9, 4]. Our titrant (CBS, Sigma-Aldrich 4-Sulfamoylbenzoic acid, product C11804, lot MKBF3323V, 97% purity by FT-NMR, MW 201.2) comes as a powder, from which we aim to prepare a solution of purity-corrected^4^ concentration [*X*]_*s*_ ∼ 720 *μ*M using the dialysate. Uncertainties in the true [X]_*s*_ come from at least two sources: the mass of CBS and volume of buffer used in preparing this solution, each of which will be imprecise due to measurement error. Further dilution steps will introduce additional error.

To load the VP-ITC syringe, we require ∼2.1 mL of our titrant^5^. For our chosen [X]_*s*_ ∼ 720 *μ*M, this requires only 0.3 mg of CBS, but given the precision of the analytical balance used for this step (Mettler-Toledo AB204, readability ±0.1 mg), this would yield 33% uncertainty in [X]_*s*_, and hence the final relative errors in *K*_*a*_ and Δ*H*° would be *at least* this large. To reduce the mass uncertainty to 1%, we must weigh out at least 10 mg. Since the solubility of CBS in water is only 453 mg/L at room temperature (which corresponds to a 2 250 *μ*M solution), we need a volume of at least 22 mL to dissolve 10 mg. Using a 25 mL Class A volumetric flask or pipette (rated ±0.05 mL) would allow us to attain the desired 1% precision. On the other hand, graduated cylinders and serological pipettes with 25 mL capacity often possess a precision of only ±0.5 ml, which would raise the uncertainty in [X]_*s*_ to 2%. Here, we found it convenient to employ multiple liquid transfers with a Gilson P5000 5 mL pipette, which has a stated reliability of ±0.03 mL at full capacity^6^.

We chose to prepare a 1 500 *μ*M CBS stock solution as a compromise between ensuring complete solubility of CBS (solubility 2 250 *μ*M in water) and minimizing buffer use (preparing a solution of ∼720 *μ*M directly with 10 *μ*g CBS would have doubled the quantity of buffer required). To do this, we added 10.0(1) mg CBS to 32.1(2) mL PBS dialysate and vortexed to ensure the compound was completely dissolved, yielding 32.2(2) mL of a 1.50(2) mM CBS stock solution.

To ensure sufficient ∼718.29 *μ*M titrant to allow for a ligand-into-buffer blank titration and additional experimental replications if needed, we planned to prepare 9 mL of titrant solution. This is more than necessary, as minimum of 700 *μ*L/experiment is required for the VP-ITC if the low-volume syringe loading tube is utilized. Using the Gilson P5000, we then added 4.309(12) mL CBS stock to 4.691(12) mL PBS to obtain a 717(9) *μ*M CBS titrant (1.2% RSE). Error propagation was performed automatically by the spreadsheet (Figure A.3).

While the use of volumetric glassware in principle requires all solutions and glassware to be equilibrated to the glassware calibration temperature, in practice, the contribution of thermal expansion to inaccuracies is generally insignificant. Due to the low coefficient of thermal expansion of borosilicate glass, this expansion will only introduce an error of 0.0010%/° C [12]—small enough to be negligible for our purposes. If gravimetric solution preparation (GSP)—in which the mass of both compound and solvent is used to determine the final concentration—had been used instead, the larger coefficient of thermal expansion of liquids can make a larger contribution to the error (dilute aqueous buffers have a coefficient of thermal expansion near that of water, ∼0.025%/° C), but still generally amounts to a negligible contribution to error for calorimetry even for changes of several degrees. Automated systems for gravimetric solution preparation and concentration error determination are available (such as the Mettler-Toledo Quantos), though more commonly used in industrial settings.

Alternatively, we could have determined the titrant concentration [X]_*s*_ spectrophotometrically using the known extinction coefficient of CBS at 272 nm (reported as *∊*_272 nm_ = 1.31(13)×10^3^ M^−1^ cm^−1^ [3]). However, since the uncertainty in the absorbance measurement is 1%, the uncertainty in the extinction coefficient *∊* (10%) would dominate the concentration error, resulting in a spectrophotometrically-determined concentration that is uncertain by ∼10%. Indeed, the concentration we measure in this manner—700(70) *μ*M—is consistent with that determined by mass and volume, but is an order of magnitude more uncertain; had we chosen to use this spectrophotometrically-determined concentration for [X]_*s*_, our final uncertainties in *K*_*a*_ and Δ*H*° would be at least 10%.

## Data analysis

The titration dataset (Figure 1) was analyzed using Origin 7.0 (OriginLab Corp.) after subtracting heats obtained from a separate ligand-into-buffer blank titration utilizing the same protocol (Supplementary Figure A.2). Here, the blank heats were small and uniform, of the same magnitude as water-into-water injections. The least-squares (LS) fit of the thermodynamic parameters to the integrated injection heats are shown in the caption of Figure 1. Note that, since the stoichiometry is known to be 1:1, the site parameter *n* absorbs errors in [M]_0_ and the cell volume V_0_; if the actual concentration of active macromolecule is of interest, these quantities will require more precision [13]. While Δ*H*° is rather insensitive to errors in the stated cell volume V_0_ as a result, those errors can have a substantial effect on *K*_*a*_, so careful calibration of V_0_ using standard reactions (e.g. [2, 10]) is advised if highly accurate *K*_*a*_ is sought [13]. Raw and processed datasets are provided with this work as Supplementary Material.

**Figure 1:**
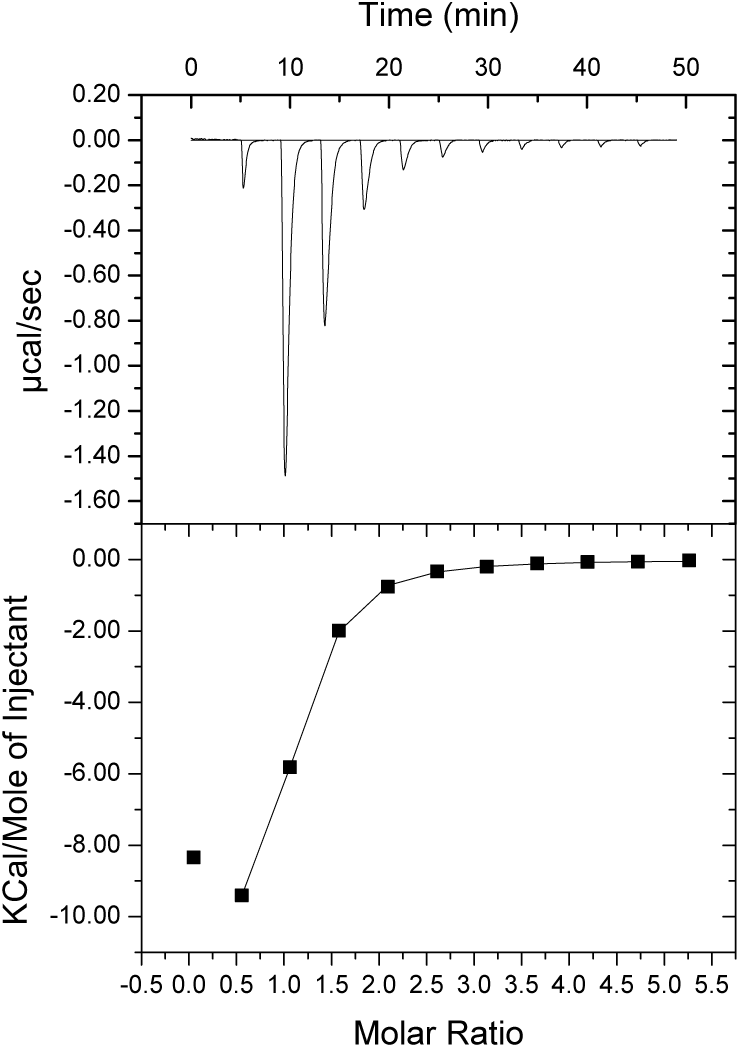
Titration of CAII by CBS at 25 C as visualized by Origin. *Top:* Differential power *vs.* time after blank subtraction and baseline correction in Origin. *Bottom:* Peak integration and 1:1 binding model fit in Origin. Least-squares fit with Origin (excluding the first injection) gives the following thermodynamic parameters and fit uncertainties: *K*_*a*_ = 1.20(3) × 10^6^ M^−1^, Δ*H* = −11.27(6) kcal/mol, *n* = 0.915(3). Propagation of titrant error with the provided spreadsheet gives updated parameters with realistic uncertainties: *K*_*a*_ = 1.20(3) × 10^6^ M^−1^, Δ*H* = −11.3(2) kcal/mol, and *n* = 0.915(4). See Supplementary Material for complete experimental details and links to download raw data.

The error reported by the LS fit only represents the error in model fitting assuming the specified concentration for titrant is *exact*—we must now include the uncertainty in the titrant concentration to obtain an estimate of the true error. Provided the relative errors in concentration [*X*]_*s*_ are sufficiently small (< 10%) for the standard Taylor expansion propagation of error above to be accurate, we can use Eq. 2 to estimate the relative error in the thermodynamic parameters *K*_*a*_ and Δ*H*° and site parameter *n* given corresponding uncertainties from the least-squares fit (*s*_K,LS_, *s*_ΔH,LS_, *s*_n,LS_) and in the titrant concentration *s*_[X]s_ [4],

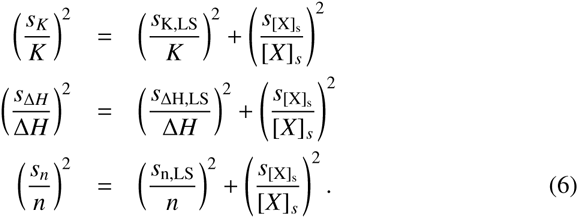

where the terms *i* and *j* from Eq. 2 are determined from the manner in which [X]_*s*_ influences the thermodynamic parameter of interest (see, e.g. Table 1 of [9]). Since the uncertainty in our [X]_*s*_ is only 1%, the 3% LS fit uncertainty dominates for *K*_*a*_; but for Δ*H*° the titrant uncertainty is more important, increasing the RSE from 0.7% to 1.2%. These computations are automatically handled by the spreadsheet, which also computes Δ*G*° and Δ*S*° and their uncertainties.

Since Δ*G*° logarithmically depends on *K*_*a*_ through the relation Δ*G*° = −*RT* ln[*K*_*a*_*C*_°_], the uncertainty in Δ*G*° computed using Eq. 6—where *s*_Δ*G*°_ = *RT* (*s*_*K*_*a* /*K*_*a*_) = 0.02 kcal/mol—is much smaller than that in Δ*H*° (0.15 kcal/mol). If the entropic contribution to binding, −*T* Δ*S*° = Δ*G*° Δ*H*° is of interest, its uncertainty can similarly be obtained from Eq. 1, and found to be of the same magnitude as that in Δ*H*° (0.15 kcal/mol)^7^.

Comparing our results including final uncertainties propagated by the spreadsheet [*K* = 1.20(3) × 10^−6^ M^−1^ and Δ*H* = −11.3(2) kcal/mol] with the best-fit to the ABRFMIRG’02 results [*K* = 1.08(4) × 10^6^ M^−1^ and Δ*H* = −11.11(4) kcal/mol] [4], we see that the difference in *K* = 0.12(5) × 10^6^ and Δ*H*° = 0.2(2) kcal/mol. The RSEs of our results are 3% in *K*_*a*_ and 1% in Δ*H*°—in line with the predicted errors from our initial experimental modeling step.

## Discussion

Note that our excess uncertainty comes directly from the uncertainty in the prepared titrant concentration [*X*]_*s*_. Had we chosen to use much less than 10 mg of compound, or utilized low-precision volume transfer devices (such as serological pipettes), we could have easily raised this contribution to 10% or more, which would then dominate our apparent LS uncertainties. Although the absolute error in Δ*G*° would remain small (∼ 0.04 kcal/mol), the absolute error in Δ*H*° would be large (∼ 1.1 kcal/mol), making the error in −*T* Δ*S*° comparable in magnitude. This can have important consequences in trying to ascribe significance to differences in entropy-enthalpy compensation within a congeneric series, especially when differences in Δ*G*° are small [14, 5, 6, 15].

We recall that the reported errors in Δ*H*° (and hence *T* Δ*S*°) for the ABRF-MIRG’02 study were as much as two orders of magnitude smaller than the actual error deduced from variation among independent measurements. If indeed concentration errors were at fault, simply repeating the experiment with the same solutions would not have revealed any problem [4, 15].

## Ackowledgments

JDC and SEB acknowledge funding from the Memorial Sloan Kettering Cancer Center, as well as from the California Institute for Quantitative Biosciences (QB3) at the University of California during earlier stages of this work. The authors are grateful to Brian K. Shoichet (UCSF) for the use of his laboratory facilities for this study, as well as Allison Doak (UCSF and Matthew Merski (UCSF) for their assistance. The authors thank Patrick Grinaway, Daniel Parton, and Ariën Sebastian Rustenburg (MSKCC) for stimulating the publication of this work through their subsequent work on this system, and Antonio Luz and Fraser Glickman (Rockefeller High-Throughput and Spectroscopy Resource Center) for their assistance and expertise with automated calorimetry.

## Appendix A. Supplementary Material

### Appendix A.1. Experimental Details

Both the CBS-into-CAII and CBS-into-buffer titrations were conducted using the following protocol in Table A.1. An archive of the MicroCal VP-ITC data files (.itc) generated by these experiments are available as Supplementary Material.

**Table A.1:** Experimental parameters for VP-ITC.

|  |  |
| --- | --- |
| initial “down syringe” following purge-refill | 10 $\mu\text{L}$ |
| number of injections | 11 |
| cell temperature | 25 C |
| equilibration time | 300 s |
| stir speed | 307 RPM |
| reference power | 10 $\mu\text{cal/s}$ |
| feedback mode/gain | high |
| equilibration | fast |
| initial “throwaway” injection volume | 1 $\mu\text{L}$ |
| subsequent injection volume | 10 $\mu\text{L}$ |
| injection rate | 2 s/$\mu\text{L}$ |
| time between injections | 240 s |
| filter time for power measurements | 2 s |

### Appendix A.2. Ligand-into-buffer titration

Supplementary Figure A.2 shows the ligand-into-buffertitration.

**Figure A.2:**
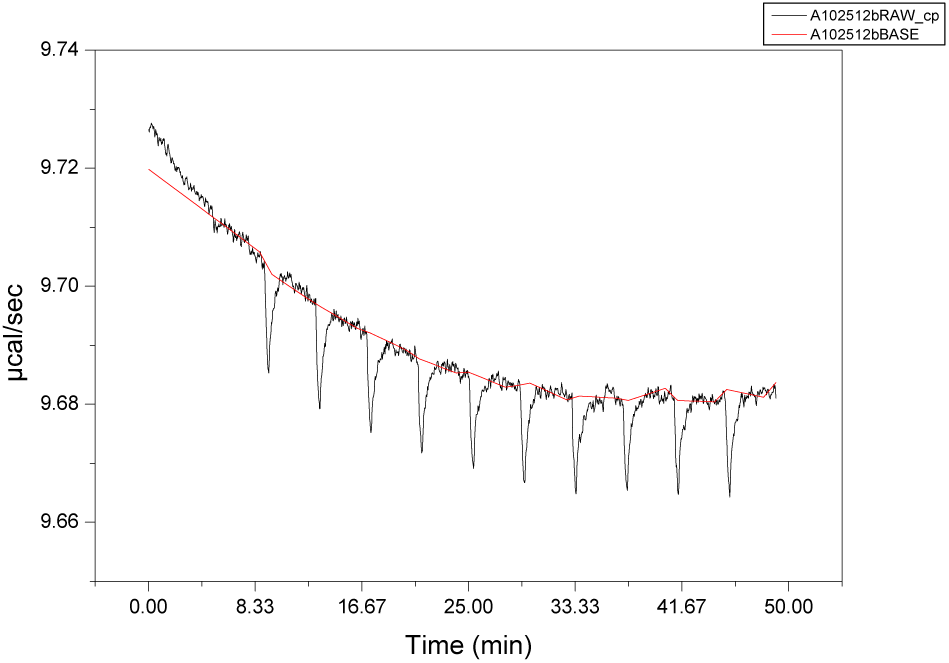
Differential power for ligand-into-buffer titration. Differential power is shown in black, with the Origin automatic baseline fit in red.

### Appendix A.3. ITC Spreadsheet

Figure A.3 depicts the spreadsheet (available for for download from Supplementary Material in multiple formats, and online at http://github.org/choderalab/itc-worksheet) with the details for the CBS-CAII titration experiment reported here filled in.

The spreadsheet is divided into sections corresponding to the different components of a typical ITC experiment. The first section (*Experimental Details*) contains general details of the experiment, the second section (*Ligand*) the details of ligand (titrant) solution preparation, the third section (*Protein*) the protein (titrand) preparation, and the final section (*Thermodynamic Parameters*) the details of the least-squares fit and overall error. Green cells indicate records the user is to fill in during the planning stages of the experiment, yellow cells are filled in by the user during the course of preparing solutions and executing the experiment, grey cells are automatically computed by the spreadsheet to aid the user in experimental design and analysis. Importantly, during both preparation of the titrant and titrand, a “typical error” sets the upper bound for the error the experimenter should be able to achieve. Exceeding this typical error is a clear indication that a precision-limiting step has crept into the workflow.

We stress that the “desired” grey fields specify target values that the experimenter is encouraged to meet as closely as possible, but the practicalities of experimental work often necessitate practical deviations from these goals. The spreadsheet is still able to allow the experimenter to track their actual measurements at each step and propagate error to the final results accordingly.

**Figure A.3:**
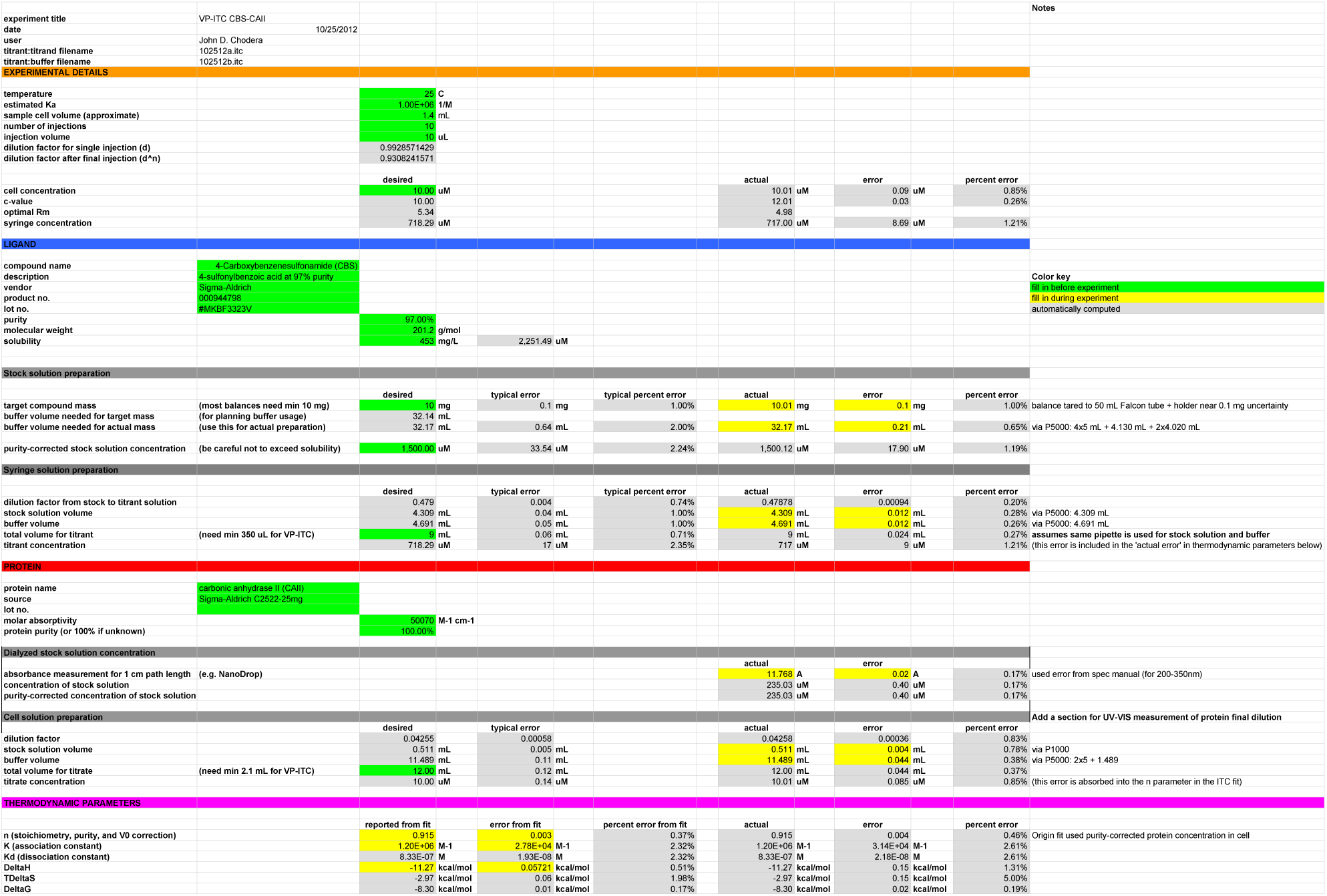
Spreadsheet for this experiment showing automated propagation of error. This spreadsheet and blank templates is available for download in multiple formats at http://github.org/choderalab/itc-worksheet. Note that some quantities are recorded to greater precision than experimental uncertainty in the spreadsheet by virtue of having been recorded directly from the instrument. These quantities are always written in the text with appropriate attention to significant figures—that is, only the largest significant figure of the uncertainty is recorded, and the value it is attached to is truncated to that decimal place.

## Footnotes

1 Here, *R* denotes the ideal gas constant, *T* the absolute temperature, and *C*_0_ the standard state concentration.

2 http://github.com/choderalab/itc-worksheet.

3 While use of this expression requires a rough estimate of the reaction *K*_*a*_ and an [*M*]_0_ that will produce observable heats, this is currently unavoidable in the practice of calorimetry. In the worst case, a pilot experiment using minimal material can be used to crudely estimate these quantities and Eq. 4 used to determine optimal conditions for a second experiment.

4 Recall that 97% purity denotes 1 g of powder should contain 0.97 g CBS.

5 A smaller 700 *μ*L filling tube is also available.

6 For pipettes, the stated systematic error Δ is generally larger than the imprecision. If the same pipette is used to deliver *multiple* aliquots, the uncertainty for the total volume transferred should be estimated from summing the systematic error Δ for each transfer. Since the systematic error is generally unknown and we utilize each pipette once, we use the stated systematic error to estimate the uncertainty in transferred volume, assuming it behaves like random error. For example, if 25 mL is transferred in five transfers of 5 mL using a P5000 (Δ = 30 *μ*L), the uncertainty is (5)(0.030) = 0.15 mL.

7 Because Δ*H*° and *K*_*a*_ (hence Δ*G*° are obtained from the same fit—and hence are correlated—cross-terms of the form 2(∂ *f* |∂*x*)( ∂ *f* |∂*y*)*s*_*xy*_ with *x* ≡ Δ*G*° and y ≡ −Δ*H*° must be added to Eq. 1, but because the uncertainty in Δ*H*° is an order of magnitude larger than that in Δ*G*°, it still dominates the overall uncertainty even if these correlation terms are included.

